# Perioral secretions enable complex social signaling in African mole-rats (genus *Fukomys*)

**DOI:** 10.1101/2022.10.04.510857

**Authors:** Kai R. Caspar, Pavel Stopka, Daniel Issel, Kristin Katschak, Till Zöllner, Sina Zupanc, Petr Žáček, Sabine Begall

**Affiliations:** Department of General Zoology, Faculty of Biology, University of Duisburg-Essen, Essen, 45147 Germany; Department of Game Management and Wildlife Biology, Faculty of Forestry and Wood Sciences, Czech University of Life Sciences, Praha, Czech Republic; Department of Zoology, Faculty of Science, Charles University, BIOCEV, Vestec, Czech Republic

**Keywords:** olfaction, sebaceous gland, sebum, Bathyergidae, odor preference

## Abstract

Subterranean common mole-rats of the genus *Fukomys* (family Bathyergidae) live in large cooperatively-breeding families. Odor cues have been hypothesized to importantly mediate social behaviors in the underground ecotope, but only little is known about the role of olfactory signaling in burrowing mammals. Here we characterize the so far neglected perioral glands of *Fukomys* and other African mole-rats as an important source of olfactory social information. Histology demonstrates these structures to be derived sebaceous glands that are developed regardless of sex and reproductive status. However, gland activity is higher in *Fukomys* males, leading to sexually dimorphic patterns of stain and clotting of the facial pelage. Behavioral assays revealed that conspecifics prefer male but not female perioral swabs over scent samples from the back fur and that male sebum causes similar attraction as anogenital scent, a known source of social information in *Fukomys*. Finally, we assessed volatile compounds in the perioral sebum of the giant mole-rat (*Fukomys mechowii*) via GCxGC-MS-based metabolomic profiling. Volatiles displayed pronounced sex-specific signatures but also allowed to differentiate between intrasexual reproductive status groups. These different lines of evidence suggest that mole-rat perioral glands provide complex odor signals that play a crucial role in social communication.

## Introduction

The strictly subterranean, tooth-digging Northern common mole-rats of the genus *Fukomys* (family Bathyergidae – African mole-rats) have become a model group to study social dynamics in cooperatively-breeding small mammals (Burda, 1990; Burland et al., 2002; Patzenhauerová et al., 2013; Zöttl et al., 2016; Torrents-Ticó et al., 2018). These sub-Saharan hystricomorph rodents live in family groups organized around a single reproductive pair that occupy and maintain extensive burrow systems (Patzenhauerová et al., 2013). Dispersal of offspring is delayed so that juveniles may stay with their parents well into adulthood, creating cohesive family groups that typically comprise around 10 members (Burda et al., 2000; Bennett & Jarvis, 2004; Torrents-Ticó et al., 2018). Social dynamics in wild *Fukomys* have been studied most intensively in the Damaraland mole-rat (*Fukomys damarensis*) of the Kalahari Desert, but are assumed to be largely uniform among congeneric species (Burda et al., 2000; Torrents-Ticó et al., 2018). Dispersal in Damaraland mole-rats is sex-biased, with males dispersing at higher rates and across longer distances than females do (Torrents-Ticó et al., 2018; Mynhardt et al., 2021). After dispersal, females will typically establish their own burrow system and will live solitarily until a mate arrives – at times for several years (Thorley et al., 2021). Dispersing males seek out solitary females, but also show a notable propensity to invade established family groups and challenge the same-sex breeder there (Young & Bennett, 2013; Torrents-Ticó et al., 2018; Mynhardt et al., 2021). This creates asymmetrical reproductive competition, which is reflected in pronounced male-biased sexual dimorphism in many *Fukomys* species (Caspar et al., 2021a).

The peculiar social system of *Fukomys* mole-rats is hypothesized to be crucially maintained by olfactory signals. In line with that, comparative genomic evidence points to excellent olfactory capacities in these animals (Stathopoulos et al., 2014). Behavioral experiments have demonstrated that group members can individually identify each other based on olfactory cues, such as anogenital scent (Heth et al., 2002). This allows the discrimination of familiar from foreign individuals and enables the strict incest taboo found among *Fukomys* families (Burda, 1995). Without regular contact to each other, however, family members will at some point cease to recognize their relatives (ca. 18 days in *F. anselli* – Burda, 1995; > 4 months in *F. mechowii* – Bappert & Burda, 2005). Interestingly, mole-rats can still differentiate such estranged siblings from total foreigners based on scent cues, which might indicate that body odors convey information about relatedness in these rodents (Heth et al., 2004). A recent study also demonstrated that *Fukomys* can distinguish between groups and single foreign conspecifics as well as identify the sex of the latter based on soil-born scents (Leedale et al., 2021). This further supports an important role of odors for social communication and implies that the search for mates in dispersing mole-rats could be guided by olfactory stimuli. A yet unappreciated source of scent signals in *Fukomys* mole-rats are their perioral secretions, which stain the cheek region adjacent to the procumbent extrabuccal incisors of the animals.

Perioral stains (“mentum” – Macholán et al., 1998) have been noted in many *Fukomys* species (*F. amatus*: Macholán et al., 1998; *F. anselli*: Begall et al., 2021; *F. damarensis*: Bennett & Jarvis, 2004; *F. darlingi*: De Graaff, 1964; *F. mechowii*: Peters, 1881; *F. micklemi*: pers. obs.; *F. vandewoestijneae*: Van Daele et al., 2013; but note that data on basal-branching species from the Northern hemisphere are missing). Typically, the stain is dark brown with a reddish to yellowish tinge and is restricted to the perioral region (Fig. 1A-D). The secretion is dry and solid, with a texture and consistency comparable to candle wax (pers. obs.). In the sister lineage to *Fukomys*, the Southern African mole-rat genus *Cryptomys*, yellow perioral stains have been reported (Fagir et al., 2021), but those are more inconspicuous than in the former (K. Finn, pers. com.). In other bathyergids, no noticeable facial stains appear to be present but it is known that many if not all rodent species exhibit perioral glands that aid in olfactory communication. These structures have been described in various taxa of hystricomorph, sciuromorph, and myomorph rodents, including the subterranean blind mole-rats of Eurasia (Quay, 1965; Sokolov, 1982), and are especially well-studied in squirrels (Brady & Armitage, 1999). Indeed, such perioral glands have already been sketchily described in the naked mole-rat (Quay, 1965; Kimani, 2013), another social bathyergid.

**Figure 1:**
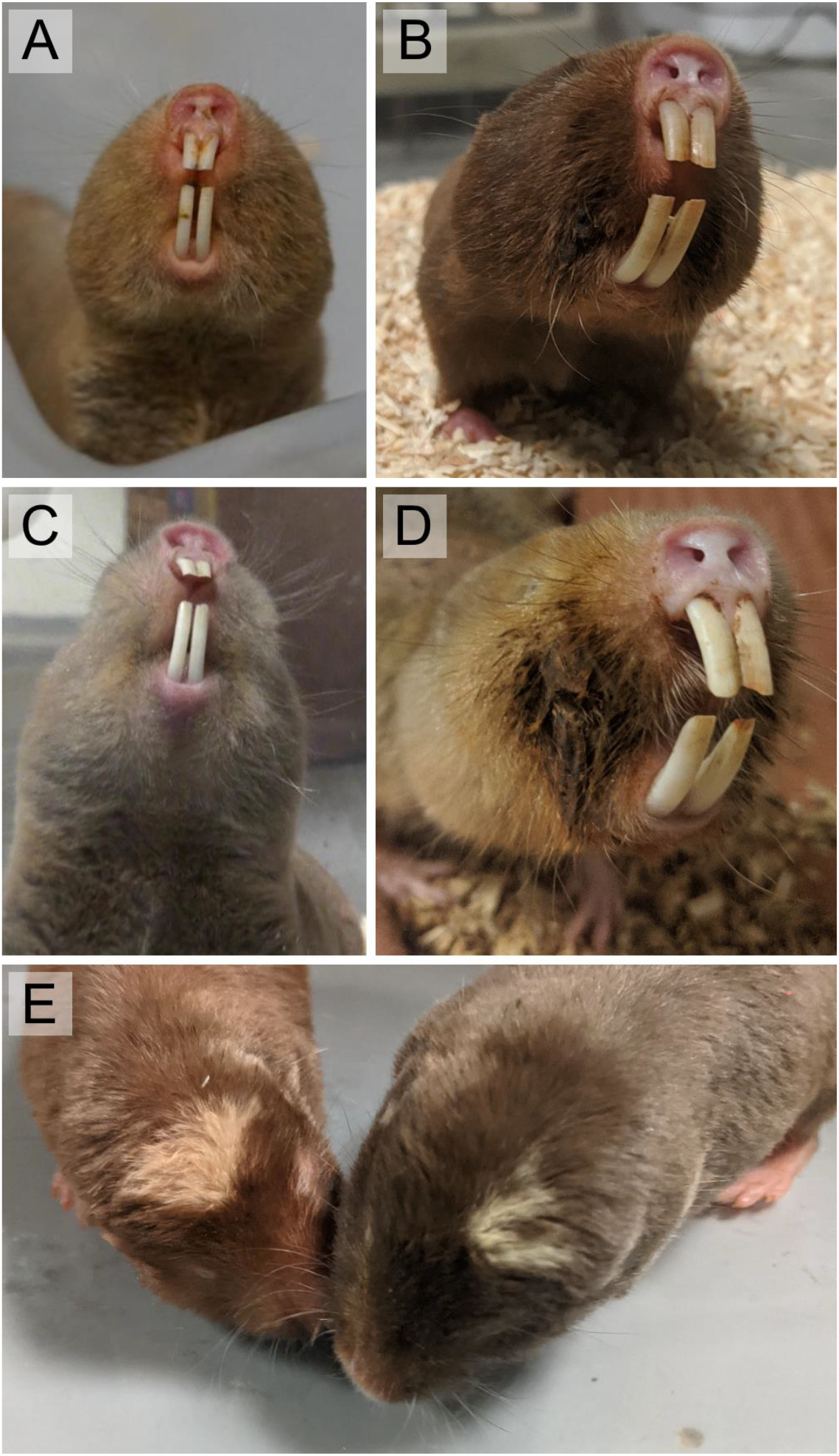
Sex-specific expression of perioral secretions in adults of different species of Northern common mole-rats (*Fukomys*) and relevant social behaviors. Note the stronger expression and clotting of the fur in males. A: Female Micklem’s mole-rat (*Fukomys micklemi*) B: Male Micklem’s molerat. C: Giant mole-rat female (*Fukomys mechowii*). D: Giant mole-rat male. The perioral secretions in this individual are particularly pronounced. E: Facial nuzzling in a freshly mated pair of Mashona mole-rats (*Fukomys darlingi*). The female (left) is sniffing the perioral region of the male.

Although perioral secretions in *Fukomys* have been known for centuries and are a striking component of the animals’ appearance (Peters, 1881) they have attracted only little attention from researchers so far and their biological significance remains enigmatic. De Graaff (1964) proposed that the stain is derived from specific food items, but as observations from the wild were accumulating and once *Fukomys* had been established in captivity, it became obvious that the stains are endogenous secretions and unrelated to general food intake.

While there is no evidence that *Fukomys* use perioral secretions to actively scent mark their environment, they could play an important role in social communication. Both *Cryptomys* and *Fukomys* engage in conspicuous reciprocal cheek nuzzling in various social situations (Fig. 1E). This behavior often initiates copulation (see e.g., Bennett, 1989 – *Cryptomys hottentotus*; Scharff et al., 1999 – *Fukomys mechowii*) but is also observed when unfamiliar individuals, regardless of sex, meet for the first time (Burda, 1989 – *Fukomys anselli*) and when family members greet each other after short periods of separation (pers. obs.). In the mating context, it typically precedes anogenital inspection (Scharff et al., 1999). Obviously, perioral scents are of interest to both partners during these interactions, suggesting a role in social and particularly sexual signaling.

Several studies hypothesized that only certain status groups of mole-rats display visible perioral secretions. In particular, it has been proposed that just reproductive males and thus one animal per family, would exhibit perioral stains. Although this idea has never explicitly been tested, various field studies report to rely this character to identify breeding males (*F. anselli*: de Bruin et al., 2012; *F. damarensis*: Mynhardt et al., 2021; Maswanganye et al., 1999; *F. mechowii*: Lövy et al., 2013). The restriction of this feature to breeding males would suggest a role in both intra- and intersexual signaling and might imply an involvement in sexual suppression of subordinate males. However, there is no consensus about whether perioral secretions are indeed specific to male breeders. For instance, Kawalika (2004) reported that in Zambian giant mole-rats (*F. mechowii*) perioral stains are displayed by both sexes irrespective of reproductive status. On the other hand, Caspar et al. (2021a) noted anecdotally that the degree of expression in perioral secretions of captive Ansell’s mole-rats (*F. anselli*) is sex-specific but non-dependent on breeding status, with males in general displaying more intense stains than females (see also Caspar et al., 2021b for *F. mechowii*). In any case, a restriction of secretion to particular status groups in mole-rat communities might have important implications for their social function.

Here, we aim to characterize the occurrence, chemical composition, and biological significance of perioral secretions in mole-rats of the genus *Fukomys* by the aid of morphological and histological observations, behavioral assays, and chemical analyses.

## Materials and Methods

All statistics were performed in R (R Core Team, 2021).

### Histology of the mouth corner integument in African mole-rats (*Fukomys* spp. & *Heterocephalus glaber*)

We sampled the perioral integument of the mouth corners in 13 *Fukomys* mole-rats, comprising three species (*F. anselli, F. mechowii*, and *F. micklemi*) as well as both sexes and intrasexual status groups (breeder vs. non-breeder), to gain insights about the morphology of glands producing perioral secretions in these animals (see Suppl. Tab. 1, for further data on sampled specimens). For comparison, we also included samples from four naked mole-rats (*Heterocephalus glaber*), another cooperatively-breeding species of bathyergid in which perioral glands but no visible secretions have been reported so far. All animals were adults deriving from the Department of General Zoology in Essen and were sacrificed for other research projects (respective animals were decapitated in deep ketamine/xylazine anaesthesia – compare Garcia Montero et al., 2015). No animals were sacrificed primarily to obtain perioral samples. Hence, the representation of species, sexes and reproductive status groups was imbalanced. Ultimately, we recovered perioral glands in the tissue samples of all of the four examined individuals of *F. mechowii* and *H. glaber*, respectively, in three out of six *F. anselli* and in none of the three *F. micklemi* (Suppl. Tab. 1). Given that we recovered glands across status groups and sexes in both genera, we are convinced that the apparent lack of glands in some individuals reflects issues with tissue sampling rather than their absence in the respective animals.

Skin was excised and prepared for standard histological sectioning and staining. *Fukomys* mouth angles were shaved with a handheld trimmer (Isis GT420; Aesculap, Suhl, Germany) before sampling. Tissue samples were fixed in 4% buffered paraformaldehyde for 24 h at 8 °C, subsequently transferred to 1 x DPBS buffer (PAN-Biotech; Aidenbach, Germany), and stored at the same temperature until being automatically dehydrated (Tissue Processor TP12 - RWW Medizintechnik; Hallerndorf, Germany) and embedded into paraffin (EG1150 H embedder – Leica Biosystems; Deer Park, USA). Embedded samples were manually sectioned (thickness: 5 µm) on a Microm HM 340 rotary microtome (Microm International; Walldorf, Germany), transferred to a digital precise water bath (Witeg WB-11; Wertheim am Main, Germany) warmed to 40 °C and subsequently mounted on glass slides. Tissue sections were stained using a standard hematoxylin-eosin manual staining protocol. ROTI®Histokitt II (Carl Roth; Karlsruhe, Germany) was used as a xylol-based cover medium for the tissue sections. Samples were examined and photographed on a VHX-600 digital light microscope (Keyence; Osaka, Japan).

We used ImageJ (Schneider et al., 2012) to take quantitative measurements of glandular cell sizes from the micrographs. We measured the area of medially sectioned mature non-pyknotic cells in perioral glands from Ansell’s mole-rats (n_males_ = 2, n_females_ = 1) and naked mole-rats (n_males_ = 1, n_females_ = 3). Subsequently, we tested whether there are sex differences as well as species differences in this variable for perioral glands. For Ansell’s mole-rats, we also checked for differences in cell size between perioral glands and ordinary, hair follicle-associated sebaceous glands. We compared cell sizes between gland types by means of the two-sided t-test and calculated Cohen’s D as a measure of effect size. To explore effects of species and sex on perioral gland cell size we computed a linear mixed effect model using the *lmer()* function from the lme4 package (Bates et al., 2015) of the following form: *log*_*10*_*(cell size) ~ species + sex* : *species*. The individual ID was additionally included as a random factor and we calculated η^2^ as a measure of a coefficient’s effect size. Normality of data as well as of model residuals was checked with the Shapiro-Wilk test.

### Occurrence of perioral staining among sexes and status groups of *Fukomys*

The degree of expression of perioral stains in two *Fukomys* species, the giant mole-rat (*F. mechowii*) and Micklem’s mole-rat (*F. micklemi*) was studied to test the influence of selected biological variables. The two species represent distantly related congeneric lineages (Ingram et al., 2004).

We examined mole-rats with monitored life histories living in the laboratories of the Department of General Zoology, University of Duisburg-Essen, Essen, and the Department of Zoology, University of South Bohemia, České Budějovice (Suppl. Tab. 2). All mole-rats were housed in social groups with food provided ad libitum.

Giant mole-rats derive from animals caught in the Zambian Ndola region and exhibit the diagnostic karyotype of 2*n* = 40. We included 30 males (14 reproductive ones, 16 non-reproductive ones) and 68 females (14 reproductive ones, 54 non-reproductive ones) of giant mole-rats, resulting in a total sample of *n* = 98. The imbalance among the sexes and the two female status groups is a result of the strongly female-biased sex-ratio of neonates found in this species (Caspar et al., 2021b), which also affected our sampling efforts for the mass spectrometric analyses (see below). The Micklem’s molerat lab lineage derives from animals caught at Kataba in Western Zambia, the type locality of the species (Chubb, 1909), and are characterized by a karyotype of 2*n* = 60. In this species, we studied 40 males (19 reproductive ones, 21 non-reproductive ones) and 32 females (20 reproductive ones, 12 non-reproductive ones), thus comprising a total sample of *n* = 72.

For documentation of the perioral stains, animals were briefly separated from their group, weighed, and photographed. Based on these photographs the degree of expression was scored on a species-specific qualitative scale from 1 (no visible secretion) to 4 (excessive secretion). Classifying criteria are enumerated in Table 1 and stain categories are visualized in Figure 2.

**Table 1:** Classification criteria for perioral stain patterns in *Fukomys mechowii* and *F. micklemi* (see also Figure 2 for visualization).

| Perioral secretion pattern | *Fukomys mechowii* | *Fukomys micklemi* |
| --- | --- | --- |
| 1 | No visible secretion | No visible secretion |
| 2 | Darkening of pelage immediate to the corners of the mouth, often with small lateral circular extensions | Darkening of pelage immediate to the corners of the mouth |
| 3 | Secretions visibly clot the pelage in the corners of the mouth and extend dorsally towards the periphery of the mystacial vibrissal field | Secretions visibly clot fur in the corners of the mouth and notably extend from them in lateral orientation. |
| 4 | Extensive wax-like secretions clotting large portions of the face and extending well into the mystacial vibrissal field | Extensive fur clotting in the corners of the mouth that extends to the periphery or into the mystacial vibrissal field. |

**Figure 2:**
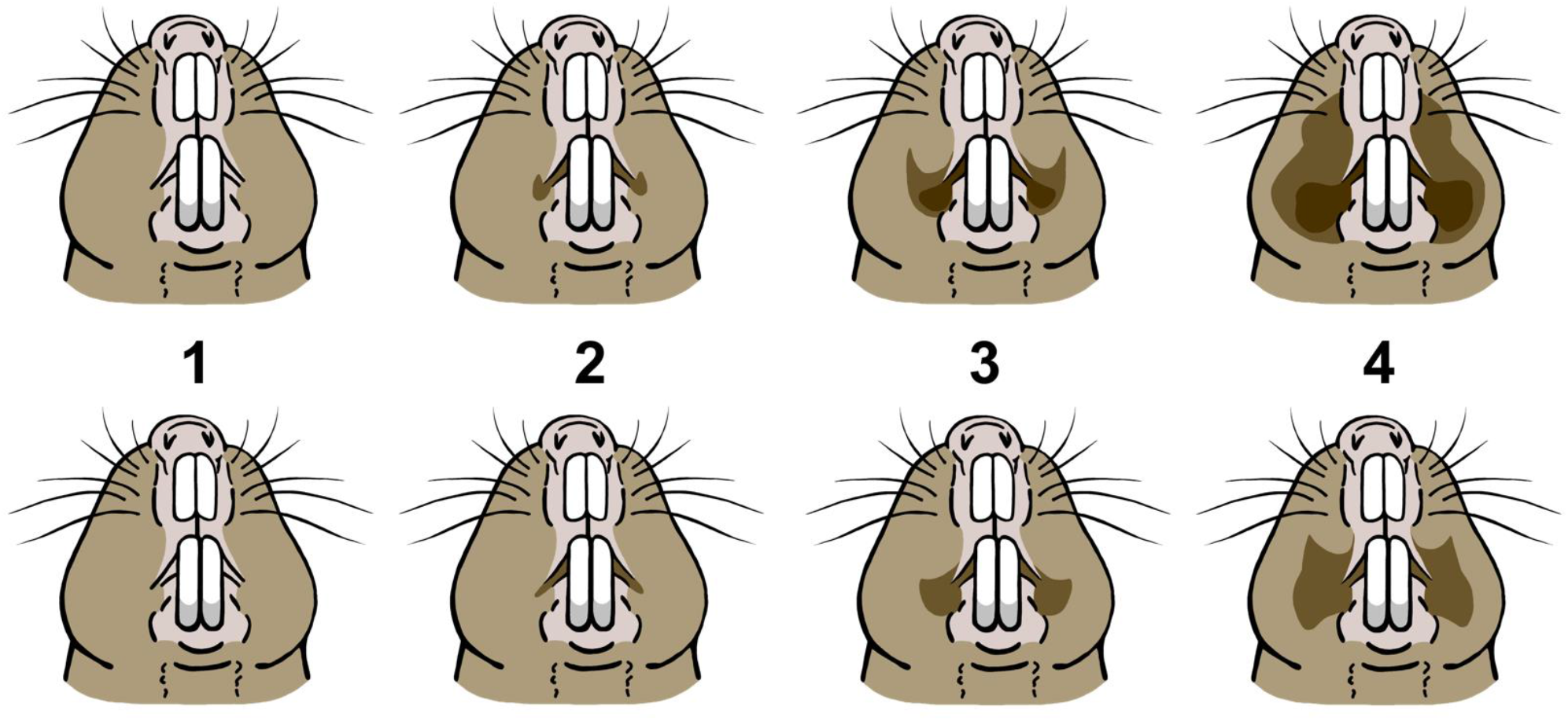
Visualization of qualitatively distinguished perioral stain patterns in *Fukomys mechowii* (top) and *Fukomys micklemi* (bottom). Compare Table 1 for a list of scoring criteria.

For each of the two species, we separately calculated cumulative link models for ordinal regression (*clm()* function of the ordinal package – Christensen, 2019; logit link function) to estimate the effects of biological variables on the expression of perioral stains. We used a two-step approach to the models: First, we used reproductive status, sex and the interactions between these two factors as model predictors to answer the question whether perioral stains are a sex-specific status signal (model I: *stain pattern ~ sex * sex* : *reproductive status*). Subsequently, we tested whether body mass (in g, log-transformed) and individual age (in months, log-transformed) predicts this trait within the sexes (model II: *stain pattern ~ sex* : *log*_*10*_*(age) + sex: log*_*10*_*(body mass)*).

### Olfactory preference tests

We ran olfactory preference assays to assess the relative informative value of mole-rat perioral secretions compared to other bodily scents. Adult (at least 17 months old) Micklem’s mole-rats (*F. micklemi*) participated in this part of the study, because the species expresses notable perioral stains while being of small size and thus easily manipulated for testing. The olfactory preference assay was designed as a two-choice set up, in which test subjects were presented with odorous swabs taken from different body regions of the same foreign conspecific donor animal with visible perioral secretions (see below). One option was invariably constituted by swabs taken from the perioral area, while the second one either derived from the donors’ dorsal pelage or perineal area. While dorsum samples acted as a simple control to simulate the presence of a foreign conspecific, anogenital smear is known to convey complex social information in *Fukomys* (Heth et al., 2002; Hagemeyer et al., 2004; Heth et al., 2004).

Odorous smear from the donor animals was collected with moistened commercial cotton swabs that were gently rubbed against the respective body region. For perioral and dorsum samples, this was done until a discoloration of the tip became visible. During the procedure, one experimenter would briefly fixate the donor animals while a second one collected the samples. The swabs were rolled out on the surface of glass cuvette lids (10.2 cm x 8.5 cm x 1.1 cm) which were subsequently used to present the odors to the test subjects (analogous to Heth et al., 2002). During this procedure as well as during the set-up of the two-choice assay, the respective experimenters wore gloves to avoid olfactory contamination of the equipment.

Test subjects were individually taken from their home cages and brought to a darkroom in which the assay set up was deployed. To allow the experimenter to operate, the room was illuminated by an LED table lamp emitting monochromatic red light (Parathom R50 80.337 E14 Red 617 nm, 6 W, Osram; Munich, Germany), which is invisible to African mole-rats (Kott et al., 2010). The animals were placed in a terrarium (50 cm x 38 cm x 30 cm) in which they were presented with the two glass plates carrying the odors of the donor animal. Glass plates were positioned equidistant from the center along the long-axis of the terrarium with a 5 cm distance to the walls and were fixed with tape on the underside to remain in place. The position (left vs. right) at which the different odor types were presented was randomized. Test animals were observed exploring the set-up for three minutes after being placed into the center of the terrarium. All experiments were recorded (SONY HDR-CX505 camcorder) and behaviors were quantified based on these recordings. Interest in the presented odors was approximated by the time spent sniffing at the respective glass plate. Sniffing was defined as the animal lowering and moving its head over the glass plate accompanied by visible movement of the rhinarium. Besides sniffing time, the latency until first sniffing for either glass plate and the number of sniffing events was quantified. However, later on these measures were deemed to be uninformative and not considered for further analyses, since the animals were for the most part alternating between the two presented options. Experimental runs in which total sniffing time was < 5 s were discarded for later analyses, leaving us with 66 valid runs in total (Suppl. Tab. 3). The researcher quantifying the sniffing responses was blinded regarding the identity of the offered scent samples.

Animals were tested in three situations. The sample sizes itemized for sex are shown in brackets:

1. *Dorsal vs. perioral secretion, male donor* (*n*_males_ = 13, *n*_females_ = 15): Mole-rats could choose between swabs from the dorsal pelage and perioral region of a foreign male conspecific.
2. *Dorsal vs. perioral secretion, female donor* (*n*_males_ = 12, *n*_females_ = 12): Mole-rats could choose between swabs from the dorsal pelage and perioral region of a foreign female conspecific.
3. *Anogenital vs. perioral secretion, male donor* (*n*_males_ = 13, *n*_females_ = 11): Mole-rats could choose between swabs from the perineum and perioral region of a foreign male conspecific. Males rather than female donors were exclusively chosen because greater interest in male secretions was found in previous runs comparing perioral and dorsal samples.

Deviations in the sample compositions for the different test situations derive from changes in the lab population caused by deaths and animals being transferred to other institutions. If possible, individual animals were tested across all three situations. A maximum of one experimental run per animal per day was performed. After each run, the set up was cleaned with water and mild detergent.

Differences in sniffing time for perioral compared to dorsum and anogenital samples were statistically assessed for each of the three test situations. Data were checked for normality using the Shapiro-Wilk test. Parametric datasets were analyzed by employing the paired t-test and calculating Cohen’s d as a measure of effect size, non-parametric ones by using the paired Wilcoxon signed rank test and Wilcoxon r to indicate effect sizes. Responses of males and females were compared for all test situations. However, sex differences were not found to be significant and thus data from males and females were pooled for all analyses to increase statistical power. Total sniffing times were compared across the three test situations by means of the Kruskal-Wallis test.

### Metabolomic profiling

We collected perioral secretions from giant mole-rats (*F. mechowii*) to identify volatile organic compounds which might act in social communication via two-dimensional comprehensive gas chromatography with mass detection (GCxGC-MS). Samples from 26 animals were analyzed (Suppl. Tab. 4; secretions from four further animals were used to calibrate the GCxGC-MS). *F. mechowii* was selected for this aspect of the study since secretions are expressed in particularly great quantities compared to congeneric species. In addition, we applied the same methodology to analyze volatiles from small amounts of hay and cereals to consider potential diet-related contamination of secretions.

Samples were collected from manually restrained live animals. Perioral secretions glue the hair in the cheek region together, so that clotted hairs could be swiftly cut and manually collected in Eppendorf tubes. Subsequently, samples were stored at -20 °C until analysis. The dynamic headspace method was used to sample the secretion compounds. The sampling process was carried out automatically using a multi-purpose sampler device (MPS, Gerstel, Germany). The sebum samples were placed in 10 ml glass vials and incubated for 5 minutes at 50 °C before a flow of nitrogen of 20 ml/min was used for continuous volatile extraction. The extraction was carried out for 10 minutes. The volatiles were sorbed on a Tenax sorbent packed in a glass tube (Tenax® TA, Gerstel, Germany) at 20 °C and subsequently released in a thermal desorption unit (TDU) at 295 °C into a programed temperature vaporizer (PTV) inlet precooled to a -30 °C where the volatiles were captured on a glass wool. The PTV inlet unit was then fast heated up to 300 °C and the analytes were introduced into a gas chromatograph. The volatiles were then analyzed employing the GCxGC-MS (Pegasus 4D, Leco Corporation, USA). A combination of non-polar and polar separation columns was used for the separation: Primary column: Rxi-5sil MS (28 m x 0.25 mm ID, Restek, Australia); Secondary column BPX-50 (1.6 m x 0.1 mm ID, SGE, Australia). Other parameters were set as follows: splitless mode, constant He flow 1 ml/min, modulation time 3 s (hot pulse 0.9 s), modulation temperature offset with respect to the secondary oven 15 °C. The temperature program applied on the primary oven: 35 °C (hold 1 min), then increase (8 °C/min) to 320 °C (hold 2 min). The temperature offset applied on the secondary column was +5 °C. Transferline temperature was held at 250 °C. The mass detector was equipped with an EI ion source and TOF analyzer enabling a unit mass resolution. The scanned mass range was 29 – 700 m/z. The ion source chamber was held at 280 °C. LECO’s ChromaTOF v4.5 was employed to control the instrument and for data processing. Selected compounds were identified by automatically matching their mass spectra with a library of mass spectra (NIST MS 2.2, USA).

To prepare the bioinformatic analysis of the GCxGC-MS data, we first generated histograms of all samples and blanks. The resulting distribution was bi-modal with compounds that occurred only in samples (green line in Figure 6A) and those that occurred in both samples and blanks (intersection). To decide which compounds are true positive metabolites, we used the mixtools package (Gentleman et al., 2004) which calculates the posterior probability (*p* < 0.05) for the identity to either of the two peaks within the mixture of two overlapping normal distributions. Next, we applied a normalization based upon quantiles, which normalizes a matrix of peak areas (i.e. intensities) with the function *normalize*.*quantiles* of the preprocessCore package (Crawley, 2007). To explore potential sources of variation in our data, we used sparse partial least squares discriminant analysis (sPLS-DA) within the mixOmics package (Rohart et al., 2017) for the fact that it has satisfying predictive performances with large datasets. To extract *p*-values of differentially abundant compounds, we used the power law global error model (PLGEM – Pavelka et al., 2004) which is an efficient tool to calculate differentiation within large data sets (e.g., proteomes, transcriptomes, metabolomes) with distributions that deviate from normality (see methods in Kuntova et al., 2018) for more details). We used ggplot2 (Wickham 2016) to visualize differentially abundant compounds.

### Ethics statement

Ethical review and approval for the behavioral assays was not required since animal housing (approved by permit no. 32-2-1180-71/328 Veterinary Office of the City of Essen) as well as all experiments complied with the corresponding animal testing regulations and were approved by the animal welfare officer in charge. No ethical permissions were necessary. All behavioral tests as well as handling protocols conformed to the relevant ethical standards and did not harm the animals. all All applied methods are reported in accordance with the ARRIVE guidelines.

## Results

### Histology of the cheek region in African mole-rats

We found large, specialized sebaceous glands in the mouth corner regions of both sexes and irrespective of reproductive status in *Fukomys* (Fig. 3A-E) as well as *Heterocephalus* (Suppl. Fig. 1). The exocrine glands in question are strongly branched, multilobated structures (Fig. 3A). Their wide secretory ducts open directly onto the skin surface (Fig. 3B; Suppl. Fig. 1). Although we were unable to measure their full extent, these glands form fields covering an area of at least several square millimeters in both species. As all sebaceous glands, they show a holocrine secretion pattern, releasing lysed cell masses into their glandular ducts (Fig. 3C). When stained using HE-solution, the glands appear well demarcated and in a light violet color. Individual gland cells are large (see below), contain well visible nuclei, and increase in size from their formation site in the peripheral layer to the center of a lobule. Perioral glands are embedded in fibrous connective tissue permeated by skeletal muscles. Otherwise, the histology of the cheek region was unremarkable and our observations aligned with those of earlier studies on bathyergid skin (Pleštilová et al., 2020; Kimani, 2013; Hesselmann, 2010; Sokolov, 1982). Regular sebaceous glands (Fig. 3F), but no other types of skin glands, were also recovered in both genera. In *Fukomys*, most hairs are arranged in follicle compounds, which are associated with one to several small, globular sebaceous glands (compare Hesselmann, 2010). In *Heterocephalus*, body hair is almost completely reduced so that regular sebaceous glands are only found in association with the vibrissae (compare Sokolov, 1982).

**Figure 3:**
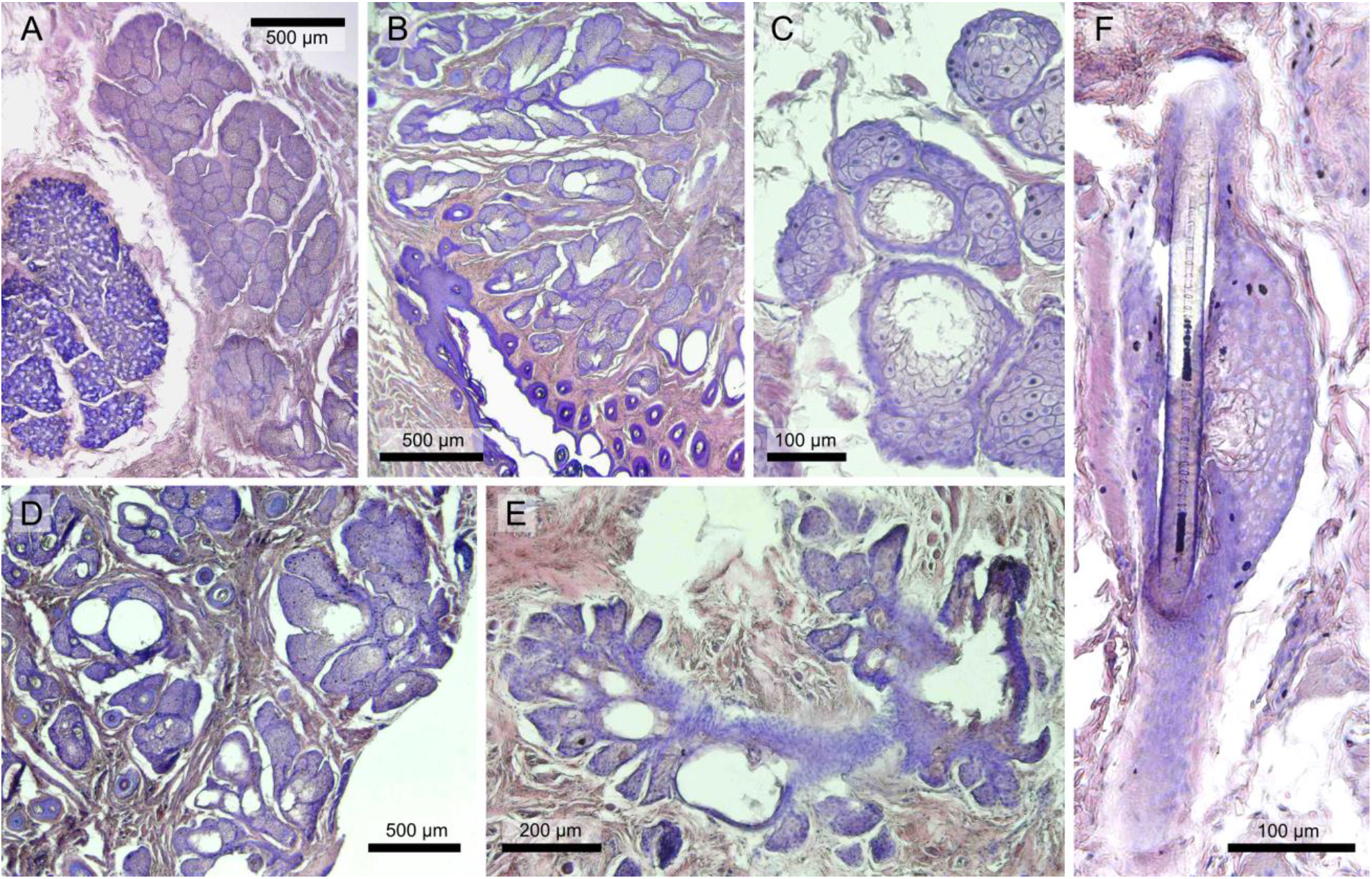
Perioral glands and regular sebaceous glands in the mouth corners of Northern common mole-rats (*Fukomys*). A: Perioral gland lobe with visible acini (right) situated deep in the dermis next to an oral mucus gland (left) in a male *Fukomys anselli*. B: Perioral gland field in a male *Fukomys anselli*. Note the wide lumina of the excretory ducts, which open directly onto the skin surface. C: Acini of perioral glands in a male *Fukomys anselli*. Note the lysis of pyknotic cells that are shed into the excretory ducts. D: Perioral gland field in a female *Fukomys anselli*. There were no obvious differences in perioral gland morphology between the sexes. E: Perioral gland in a male *Fukomys mechowii*. F: Regular sebaceous gland in *Fukomys anselli*. Note the simple globular morphology and association with a hair follicle.

Apart from deviations in morphology, cell size between perioral and regular sebaceous glands differed significantly in *Fukomys anselli*. Cells from regular sebaceous glands had a median area of 135.4 µm^2^ (*n* = 69; SD: 33.71) in the studied sections, while it was 314.7 µm^2^ (*n* = 279; SD: 92.16) for perioral gland cells (*t* = 26.105, *p* < 10^−15^; *d* = 2.23). There were no significant differences between the sexes in the size of the cells constituting ordinary sebaceous glands (*t* = 0.218, *p* = 0.831; *d* = 0.09).

Results of the linear mixed effect model on perioral gland cell sizes in bathyergids are summarized in Table 2. The regression model revealed perioral gland cell size not to differ notably between *Fukomys* and *Heterocephalus* (*t* = -2.293, *p* = 0.107; η^2^ = 0.37). However, we found a significant sex difference in cell size in *Heterocephalus* (*t* = 3.291, *p* = 0.047; η^2^ = 0.55), with the male exhibiting larger cells (*n* = 92; median: 365.5 µm^2^; SD: 113.76) than the females (*n* = 222; median: 192.1 µm^2^; SD: 72.76). In *Fukomys*, no such dimorphism was recovered (*t* = 0.111, *p* = 0.919; η^2^ < 0.01).

**Table 2:** Statistical key figures for the linear mixed effect model on perioral gland cell sizes in *Fukomys anselli* and *Heterocephalus glaber*.

| Coefficient | Estimate | Std. Error | p-value |
| --- | --- | --- | --- |
| Species (*H. glaber*) | -0.193 | 0.084 | 0.107 |
| Species (*F. ansellii*) : Sex (male) | 0.010 | 0.089 | 0.919 |
| Species (*H. glaber*) : Sex (male) | 0.277 | 0.084 | **0.047** |

### Occurrence of perioral staining among sexes and status groups of *Fukomys*

We found perioral stains to be highly sex-specific but not related to reproductive status in both studied species. Patterns of sex and status-dependent perioral stain expressions in *Fukomys* are shown in Figure 4 and are itemized in Supplementary Table 3. Males tended to show a greater development of stains than females and excessive perioral secretions (category 4) were exclusively found among the males of both species (Fig. 4). Secretion was found to be exaggerated and more strongly sexually dimorphic in *Fukomys mechowii* compared to *Fukomys micklemi* (Fig. 4).

**Table 3:** Results from ordinal regression models on perioral stain expression in two species of *Fukomys*. For each species, two models were calculated. Model I tested for effects of sex and reproductive status on perioral stain expression, while Model II did so for intrasexual effects of age and body mass.

| *Fukomys mechowii* (n = 98) |  |  |  |
| --- | --- | --- | --- |
| Model I |  |  |  |
| Coefficient | Estimate | Std. Error | p-value |
| Sex (m) | 5.745 | 1.185 | < 0.001 |
| Sex (f) : Status (repro) | -0.784 | 0.708 | 0.269 |
| Sex (m) : Status (repro) | 0.805 | 0.821 | 0.327 |
| Model II |  |  |  |
| Sex (f) : Age | -0.208 | 0.375 | 0.580 |
| Sex (m) : Age | 1.892 | 0.807 | 0.019 |
| Sex (f) : Mass | -0.153 | 0.755 | 0.840 |
| Sex (m) : Mass | -0.5187 | 0.815 | 0.524 |
| *Fukomys micklei* (n = 72) |  |  |  |
| Model I |  |  |  |
| Coefficient | Estimate | Std. Error | p-value |
| Sex (m) | 2.805 | 1.124 | 0.013 |
| Sex (f) : Status (repro) | 1.260 | 1.159 | 0.277 |
| Sex (m) : Status (repro) | 0.521 | 0.586 | 0.374 |
| Model II |  |  |  |
| Sex (f) : Age | 1.330 | 0.969 | 0.170 |
| Sex (m) : Age | 0.416 | 0.475 | 0.382 |
| Sex (f) : Mass | -0.598 | 1.476 | 0.685 |
| Sex (m) : Mass | 0.739 | 1.378 | 0.592 |

**Figure 4:**
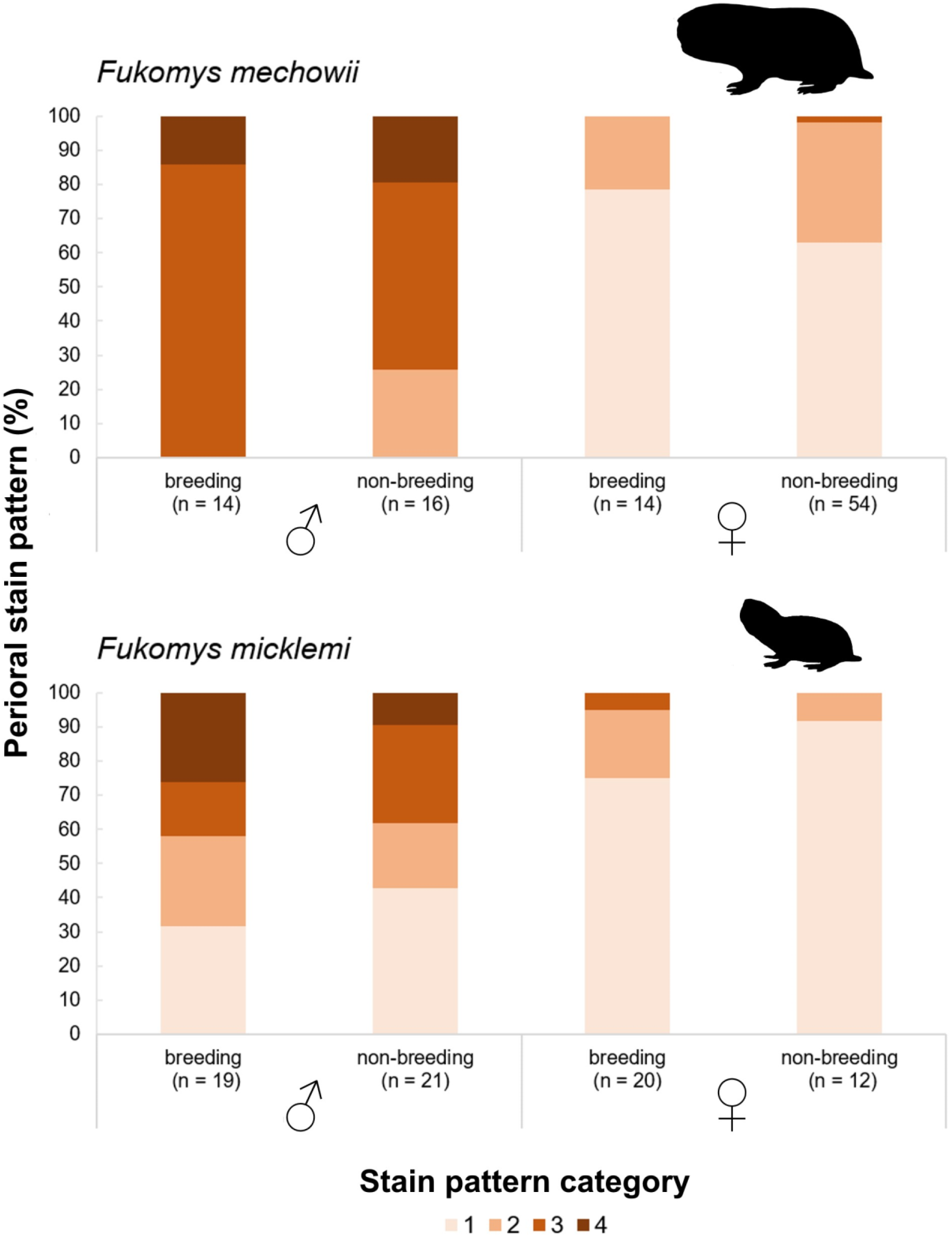
Distribution of perioral stain patterns across sexes and reproductive status groups in two species of Northern common mole-rats (*Fukomys*).

The results of the ordinal regression analyses on stain expression are provided in Table 3. In our initial models, we found that in both species sex is a good to excellent predictor of perioral stain intensity, with males showing a more pronounced expression than females (Table 3, *p* < 0.02). However, reproductive status does not influence stains in either sex or species (Table 3, *p* > 0.2). The subsequent models tested whether individual age or body mass might intrasexually influence the expression of stains. We found that these factors had no significant effects on these traits in either sex in *F. micklemi*, while we found age to be a significant predictor of stain expression in male *F. mechowii* (*p* < 0.001) exclusively (Table 3).

Note that we only sampled adult individuals here and did not systematically study when stains formed during ontogeny. Our anecdotal observations suggest that stains manifest at an age between 12 and 18 months. The youngest individual in which we noticed perioral stain was a 7-month-old *F. mechowii* female, which was sampled for the characterization of volatile compounds (Suppl. Tab. 4).

### Olfactory preference tests

Micklem’s mole-rats showed great interest in male but not female perioral swabs, although individual differences in responses were pronounced (Fig. 5). The animals showed a highly significant preference (paired Wilcoxon test; *V* = 353, *p* < 0.0001, *r* = 0.76) for odor derived from male perioral secretions (median sniffing time: 23.17 s; SD: 26.3) over swabs from the dorsum (median sniffing time: 12.45 s; SD: 14.9) irrespective of sex. There was one dropout run for this condition (final sample: *n*_males_ = 13, *n*_females_ = 14). There was a pronounced difference in the median sniffing time of female test animals (43.6 s) compared to males (14.1 s) for the perioral swabs, but it nevertheless failed to be significant (Wilcoxon test; *W* = 63, *p* = 0.19, *r* = 0.26). In fact, significant sex differences were recovered in none of the three experimental conditions.

**Figure 5:**
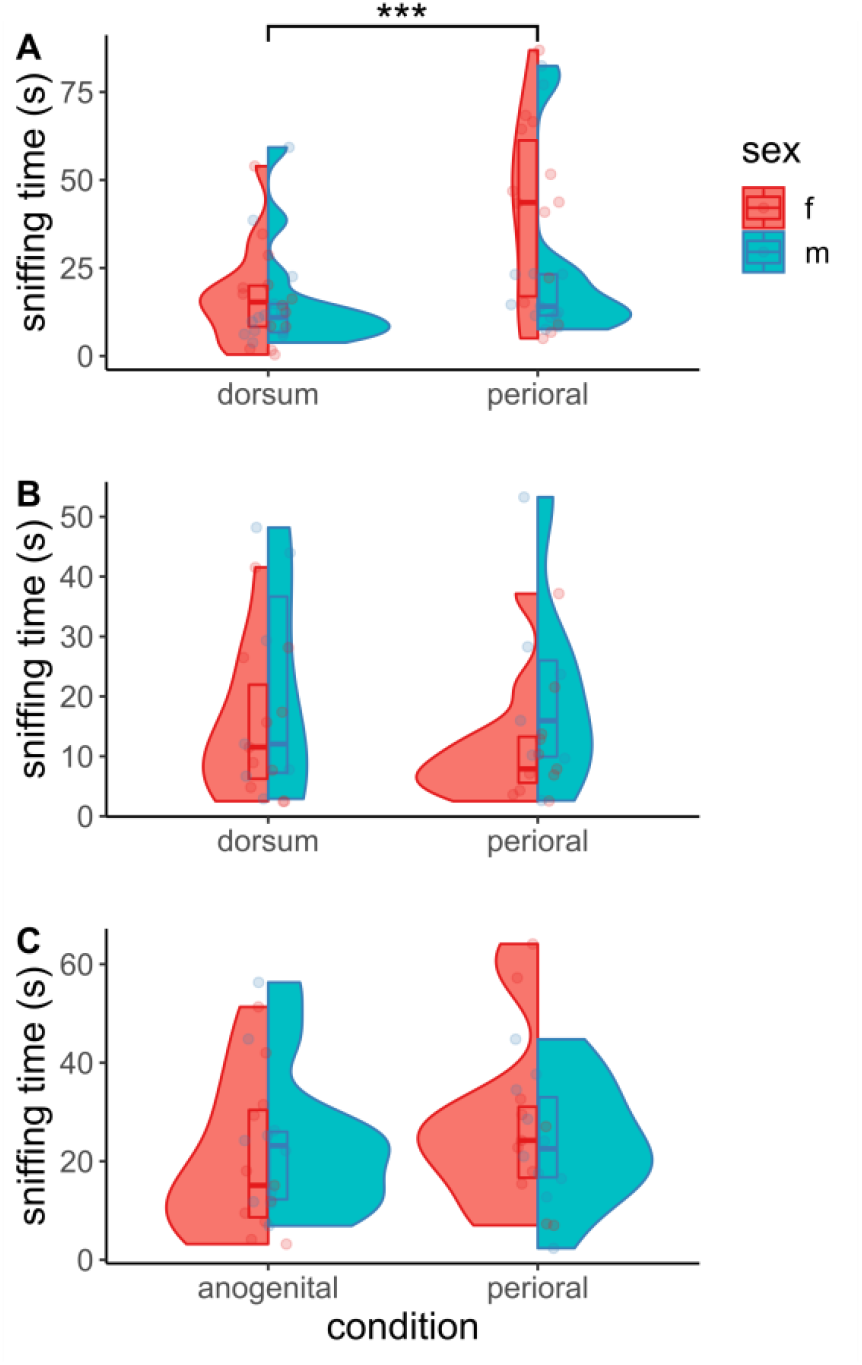
Interest of Micklem’s mole-rats (*Fukomys micklemi*) in swabs of selected body odors from foreign conspecifics. **A**: Dorsal pelage vs. mal perioral sebum. **B**: Dorsal pelage vs. female perioral sebum. **C**: Anogenital secretion vs. perioral sebum. Only condition **A** yielded significant preferences for one of the offered options.

**Figure 6:**
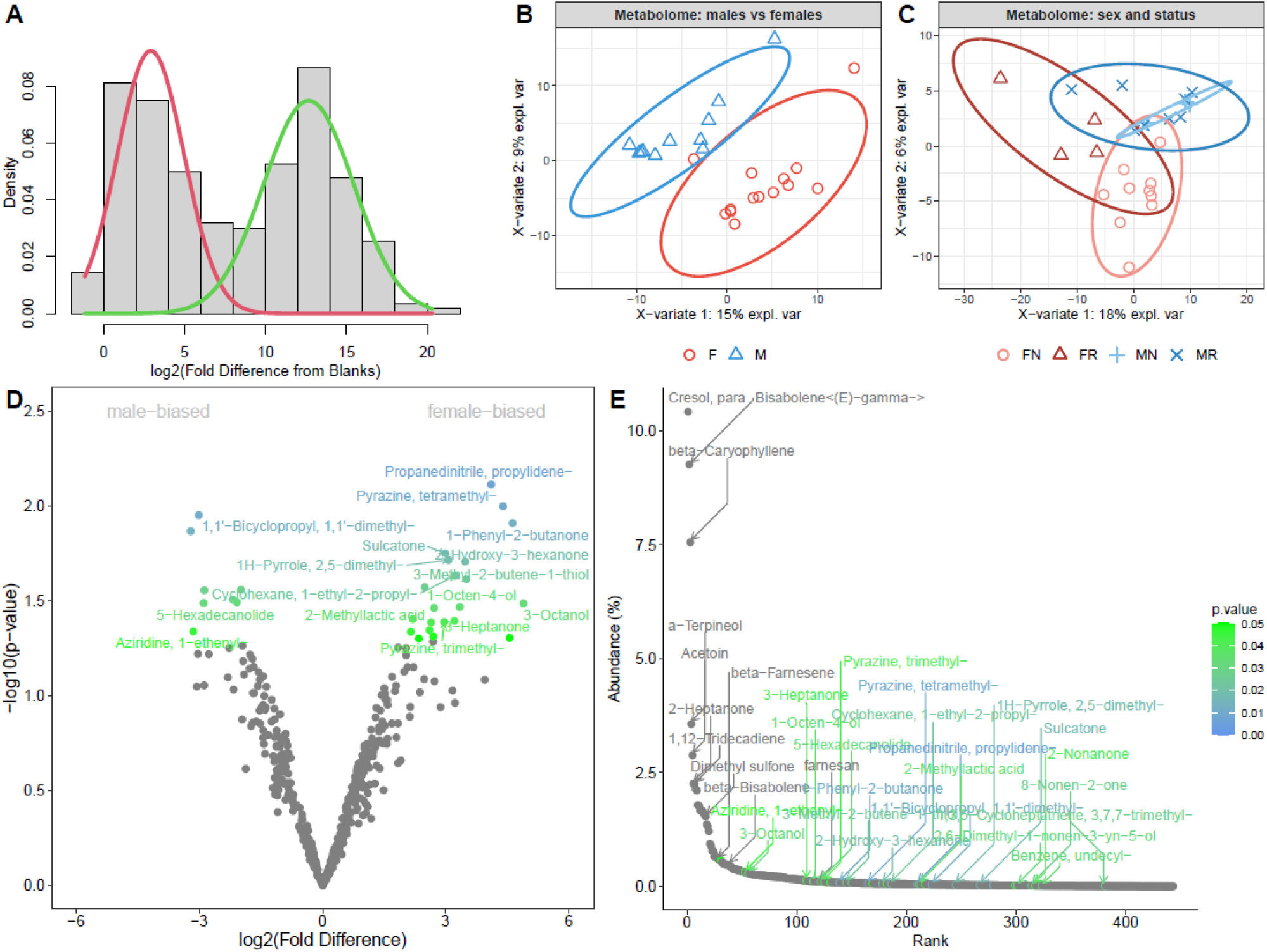
GCxGC-MS/MS analysis of volatile metabolomes from perioral secretions in *Fukomys mechowii*. **A**: The distribution of fold differences between samples and blanks is binomial while Gaussian modelling served to separate true (green line) from false positives. **B**: Sparse partial least-squares discriminant analysis revealed perfect discrimination between males (M) and females (F) and similarly reproductive status (R: reproductive, N: non-reproductive) is detectable in individuals of either sex (**C**). **D**: Volcano plot showing the distribution of female-biased and male-biased compounds. **E**: Abundance plot illustrating that most abundant volatiles are less likely to be sexually dimorphic. Colors are scaled from green (*p* < 0.05) to blue in **D** and **E**.

In contrast to male-derived samples, interest in female perioral swabs (median sniffing time: 10.25 s; SD: 13.0) was not different from that for swabs taken from the dorsal pelage (median sniffing time: 11.79 s; SD: 14.6) of donor animals (Wilcoxon test; *V* = 58, *p* = 0.39, *r* = 0.21). Additionally, we noticed the largest number of dropouts for this condition, with 6 out of the total 10 dropouts being observed here. This left us with 18 valid runs (*n*_males_ = 7, *n*_females_ = 11). Given this sex-specific difference, we continued with testing preferences for male perioral secretions compared to anogenital smear, which is a known source of social information in mole-rats. There was no significant difference in sniffing times between male perioral swabs (median sniffing time: 24.1 s; SD: 15.24) and the perineal swabs (median sniffing time: 18.05 s; SD: 15.19) examined by the mole-rats (t-Test; *t* = 1.38, *p* = 0.18, *d* = 0.24). There were three dropout runs in this condition (final sample: *n*_males_ = 10, *n*_females_ = 11).

The Kruskal-Wallis test did not indicate significant differences in the total time spent sniffing at both the two odor samples, across the three test situations (Kruskal-Wallis *?*^2^ = 3.93, *p* = 0.14).

### Metabolomic profiling

GCxGC-MS profiling of perioral secretions in *Fukomys mechowii* yielded a total of 765 volatile metabolites. However, some metabolites were also partially present in blanks. So, we employed Gaussian modeling to extract posterior *p*-values and used only those samples that were significantly not-belonging to blanks or to a group of false positives, thus yielding a total of 443 ‘true’ positive samples (all data under the green Gaussian curve in Fig. 6A; Suppl. Tab. 5).

To explore whether GCxGC-MS profiles reflect sex, we employed a supervised form of discriminant analysis, sPLS-DA, that relies on the class membership of each observation. The sPLS-DA model, based on significant components, accounted for 15 % (component 1) and 9 % (component 2) of the data variance (Fig. 6B). We used the Area Under the Curve (AUC) to provide evidence that the discrimination is perfect in both dimensions (AUC1 vs. AUC2) thus yielding AUC = 0.916 and *p* = 0.0004 for component 1, and AUC = 1, *p* = 0.00002 for component 2. This analysis thus indicates a strong sexual dimorphism in perioral sebum volatiles. Our AUC approach also revealed that component 2 is more informative than comp. 1 when looking at separations based on the reproductive status within each sex (Fig. 6C). Female non-breeders are most different from the remaining status groups (AUC = 0.99, *p* = 0.0001) but all others could also be reliably differentiated (female breeders – AUC = 0.96, *p* = 0.004; male breeders – AUC = 0.88, *p* = 0.0025; male non-breeders – AUC = 0.88, *p* = 0.037). It should be pointed out that male non-breeders cluster completely within the odor space of breeders, while the differentiation among female status groups appears far more pronounced (Fig. 6C). Yet, each separation is significant on comp. 2 and thus volatile profiles may have the potential to signal reproductive status.

To further test the hypothesis that males and females have different profiles of volatiles we used PLGEM models of differential expression/abundance to extract levels of sexual dimorphism. A volcano plot (Fig. 6D) visualizes the striking differences between the sexes, which are already detectable at the bottom of this highly symmetrical plot. However, strictly statistically speaking, when fold difference is set to 2 and *p*-value to *p* < 0.05, a total of just 28 compounds are sexually dimorphic with 7 compounds being male-biased and 21 ones being female-biased. Next, we asked whether those compounds that are sexually dimorphic are also highly abundant. Thus, we recalculated signal intensities to abundances (0 - 100%). In Figure 6E, we clearly see that those compounds that are most abundant (> 0.29%) are least likely to be sex-biased, or in other words sexual dimorphism is expressed by many compounds with rather smaller abundances while the species-specific odor-space comprises non-dimorphic compounds which are highly abundant.

The comparison of volatiles between perioral sebum, hay, and cereals demonstrated a similar number of identified compounds in the food items and the secretions (Suppl. Fig. 2; Suppl. Tab. 6). Yet only ~10 % (*n* = 68) of the total compound diversity was shared between all the three sets and even fewer being exclusively shared between sebum and hay (*n* = 2) as well as sebum and cereals (*n* = 3). We can thus exclude notable biases due to food intake for our chemical analysis.

## Discussion

### Histology of perioral glands

This study is the first to report the presence and histology of enlarged and complex perioral glands in male as well as female *Fukomys*. For *Heterocephalus*, they had been previously discussed in passing by Quay (1965) and Kimani (2013). The anatomy of the perioral glands complies to a pattern reported from all hystricomorph rodent taxa studied so far (*Capromys pilorides, Hystrix indica, Myocastor coypus* – Quay, 1965; Sokolov, 1982), in that exclusively sebaceous glands and no sudoriferous components constitute the structures. However, to our best knowledge, secretions of perioral glands in all these taxa, or in fact any other rodent, do not permanently clot the fur in a way similar to *Fukomys*, suggesting their condition to be exceptional. Interestingly, the mouth corners are devoid of comparable secretions in *Heterocephalus*.

The finding of sexually dimorphic perioral gland cell sizes in *Heterocephalus* was surprising, as this species displays remarkable monomorphism in other morphological and physiological traits (e.g., Jarvis 1991). Sex differences in gland cell size could indicate that specific secretion variables might deviate between male and female *Heterocephalus*. Yet, similar differences are not evident in *Fukomys*, although gland activity is markedly higher in males than in females of this genus (Fig. 4, see below). In any case, our findings must be interpreted with caution, as the number of sampled individuals is low for both genera and only includes a single *Heterocephalus* male as well as one *Fukomys* female. Although *Heterocephalus* has been extensively studied in regard to its social organization and communication (Buffenstein et al., 2021), chemical signaling in these animals remains essentially unknown, so that perioral secretions might represent a promising subject for future research.

Apart from the conspicuous perioral glands we describe here, no integumental scent glands are documented in *Fukomys* so far. However, it is known that various hystricomorph rodents, including *Heterocephalus*, possess specialized sebaceous skin glands in the anal area (Sokolov, 1982; Kimani, 2013). Tullberg (1899) even described such anal glands in the Southern common mole-rats of the genus *Cryptomys* (= “*Georychus coecutiens*”), providing additional indication for their presence in *Fukomys*. Secretions from these glands might well underlie the great social significance of anogenital scents in *Fukomys* (Heth et al., 2002, 2004) and can be expected to complement olfactory signaling via perioral secretions.

### Occurrence of perioral staining among sexes and status groups of *Fukomys*

We found perioral stains to be strongly sexually dimorphic but not affected by reproductive status in either of the two studied *Fukomys* species. Thus, although both sexes possess well developed perioral glands, the quantity of secretion is typically far greater in males. This might suggest that sexual hormones affect perioral secretion patterns. Indeed, it has been shown that the activity of regular sebaceous glands in rodents is stimulated by androgens and inhibited by estrogens (Thody & Shuster, 1989). Androgens have also been demonstrated to increase the size and activity of the supracaudal sebaceous gland of the guinea pig (*Cavia porcellus*), another hystricomorph species (Martan, 1962). The specialized perioral sebaceous glands of *Fukomys* might similarly respond to these hormones, giving rise to the observed sex differences in fur stain at the mouth corners.

Our analyses demonstrate that reproductive status does not notably influence the expression of perioral stain in adults of either sex. Hence, we challenge the assumption that this is a characteristic trait of breeding males (Maswanganye et al., 1999; de Bruin et al., 2012; Lövy et al., 2013; Mynhardt et al., 2021). But if adult male breeders and non-breeders essentially display equally noticeable perioral secretions, why do field studies often report it from just one animal per colony? A simple explanation might lay in the dispersal behavior of wild *Fukomys*. At the time when perioral secretions start to be notably developed, which appears to be at an age between 12 and 18 months, many male non-breeders have already dispersed from their natal family. Long-term field studies on *F. damarensis* indicate that male dispersal happens at a mean age of 12 months already, thus limiting the time that a non-breeder with fully developed perioral secretions might be captured in its natal colony (Torrents-Ticó et al., 2018). However, the time of dispersal is highly variable and it is thus not unlikely that fully adult sons may be captured along with their fathers when multiple groups are sampled. This complicates the identification of the breeding male based on perioral secretions alone and calls for a careful diagnosis of breeding status that takes other traits into account. Besides perioral stain, various studies report that male breeding status can be assessed by palpation of the testes, which are assumed to be larger in breeding males (e.g., Mynhardt et al., 2021). However, available data on whether relative testes mass is greater in reproductive compared to non-reproductive *Fukomys* males are conflicting (de Bruin et al., 2012; Garcia Montero et al., 2016).

In case of doubt, a promising indicator might instead be the width of the upper incisors. Observations on captive mole-rat families suggest that incisor width is a good relative age marker, particularly in males (Burda 2022; pers. obs.). For Zaisan mole voles (*Ellobius tancrei*), a subterranean murid species with bathyergid-like extrabuccal incisors, it has already been demonstrated that incisor width can act as an age marker well into adulthood in both sexes, before values for different age cohorts will eventually converge (Kuprina & Smorkatcheva, 2019). At what age incisor width becomes uninformative to differentiate breeding males from younger non-breeders in *Fukomys* remains to be determined.

### Olfactory preference tests

As indicated by non-significant differences in total sniffing time across experimental conditions, the mole-rats’ interest in foreign conspecific odors was comparable across the three test situations. However, dependent on the offered scents, subjects would allocate the time spent sniffing to one of the two options. Male perioral secretions were found to be preferred by conspecifics over scent samples taken from the same individual’s dorsum and to exhibit a comparable attractiveness to anogenital odor. Anogenital smear evidently conveys rich social information in *Fukomys* (Heth et al., 2002; Heth et al., 2004) and exceeds other body scents in its quality to effectively signal individual identity (Hagemeyer et al., 2004). The equivalence of perioral and anogenital swabs in the preference assay thus suggests an important communicative role for male perioral secretions in *Fukomys*. For female perioral swabs, we did not find a significant preference over samples taken from the back pelage. This could indicate a difference in the perceived informative value of female perioral odor or might simply be the result of less secretions being produced by females, resulting in a fainter scent. A possible biasing factor for our behavioral assays might be that we have not considered the reproductive status of neither the donor nor the test subjects although our subsequently generated results on perioral metabolomics suggest that mole-rats could be able to differentiate these status groups based on sebum volatiles and perhaps adopt their behavior accordingly (see below). However, as we quantified the relative informative value of odor sample pairs derived from the same respective individuals, we would not expect this issue to be a notable confounding factor here (see also Bappert et al., 2012). Nevertheless, the reproductive status of scent-sampled individuals should definitely be considered in future studies on these animals.

In any case, these results from the assays further indicate an asymmetric signaling function for perioral odors, with males investing more in the quantity of secretions to convey socially relevant signals to conspecific receivers than females do to evoke more notable responses. The observation that females spent notably longer examining male secretions than the opposite sex did, might suggest that male perioral sebum has a particular role in intersexual communication.

### Metabolomic profiling

Our analyses of volatile compounds detected via GCxGC-MS demonstrate notable individual variation and striking sexual dimorphism in the volatile chemical composition of perioral secretions in *Fukomys*, while they provide additional evidence for reproductive status-dependent signaling in both sexes. However, greater sample sizes are needed to robustly confirm this pattern and to confidently identify compounds that differentiate between intrasexual reproductive status groups, particularly male ones. Although sexual signatures were highly significant, it should be pointed out that the majority of compounds, in particular the ones occurring in the highest concentrations, are found among both sexes and can thus be expected to signal species identity. Highly sex-specific compounds represented only a fraction of the diversity and quantity of detected volatiles. Therefore, the proportional mixture of several compounds conveys a sexual signal in *Fukomys* perioral secretions, a pattern also known from the scent glands of various other rodents (Schulte et al., 1994).

Our GCxGC-MS approach used a method which compares mass spectra and retention indices of the detected compounds from perioral secretions to those in an existing library to identify volatiles. Compound names are thus just the most likely estimates. However, some of these compounds have been intensively studied and represent ‘good matches’ even without using external standards. Many of the metabolites that we detected were previously characterized in various other organisms including bacteria, plants and other animal species. This suggests that important fractions of the recovered compound diversity are not of endogenous origin but are produced by microbes colonizing the perioral sebum. This aligns well with the finding that sebaceous secretions in other mammals also house rich microbiota (compare Leclaire et al., 2014). The notably small overlap in compounds between sampled food items and sebum demonstrates that food matter did not noteworthy bias our analyses. Nevertheless, besides compounds of endogenous and microbial origin, odorants presented in the perioral area might also derive from yet other sources. For instance, the sebum might act as a hydrophobic sponge that adsorbs additional odoriferous compounds from urine or feces that are transferred to the perioral region during (auto)coprophagy or anogenital autogrooming. Future studies should aim to clarify the exact origins of odorous compounds from the perioral sebum and what information mole-rats can deduct from them. The most abundant volatile in both sexes was para-Cresol which conveys a typical ‘pig smell’ and is also secreted by male elephants during musth (Rasmussen & Perrin 1999). We also detected two variants of Bisabolenes (Bisabolene<(E)-gamma- > and beta) to be highly abundant. These sesquiterpenes are produced by many plants as well as by fungi and also function as pheromones in insects (Aldrich et al., 1993). Caryophyllene (3rd most abundant compound) is a natural sesqui-terpene present for example in cannabis and hops which has a high affinity to the CB2 receptor in mice, with strong anti-inflammatory effects (Alberti et al., 2017). Similarly, 2-Heptanone is a ketone that stimulates alarm reactions in insects, while it evokes anxiety reactions in mice and rats, even without involvement of the vomeronasal organ (Gutiérrez-García et al., 2018). This compound was also found to be abundant in both sexes.

Hence, it is clear that multiple sources contribute to the general mole-rat perioral odor space. This pattern was mirrored by sex-biased metabolites. For example, tetramethyl-Pyrazine is a bacterial metabolite and is significantly female-biased in our data. Likewise, 1-Phenyl-2-butanone is significantly female-biased, a compound that was previously detected in defensive secretions of various invertebrates including millipedes (Makarov et al., 2010). Sulcatone is also female-biased and represents a ubiquitous eukaryote metabolite. 5-Hexadecanolide is significantly male-biased in our data and, interestingly, is known to act as a pheromone in the queens of the Oriental hornet (*Vespa orientalis* – Raina & Singh 1996). 1-Ethenylaziridine is also male-biased and has previously been detected in various bacterial species (Filipiak et al., 2012).

An influence of reproductive and/or social status on volatile compounds of sebaceous gland secretions, as we observed in giant mole-rats, has also been demonstrated in a number of other social mammals, including rodents (Pohorecky et al., 2008; Zidat, 2018), primates (Setchell et al., 2010), and carnivorans (Leclaire et al., 2014). Respective odor profiles have been hypothesized to provide an honest signal of rank and physiological condition to potential competitors (Setchell et al., 2010; Zidat, 2018) and thus might aid in reducing tension and aggression. Sex and status-dependent signals from the perioral glands might serve this role in *Fukomys* families as well, facilitating the identification and social evaluation of both group members and foreign individuals that might enter an established family (compare Bappert et al., 2012).

Whether group-specific differences are adaptive or not, they might proximately be determined by the hormonal status of a respective individual. It has been demonstrated that sex hormone levels as well as pregnancy and lactation can affect commensal microbial communities and do regulate body odor via this path (e.g., Setchell et al., 2010; Pohorecky et al., 2008; Kean et al., 2011). To which extent these factors might explain differentiation in *Fukomys* perioral volatile diversity remains to be determined.

Given that perioral nuzzling is typically initiating copulation (e.g., Scharff et al., 1999), it might be tempting to speculate that certain volatiles act as sex pheromones in mole-rats. Indeed, some of the compounds we detected have been suggested to represent sexual pheromones in mice, for instance 2-Heptanone (Thoss et al., 2019). However, contrary to expectation, these molecules show no sexually dimorphic expression pattern in giant mole-rats. Apart from that, it should be noted that the vomeronasal system, which typically responds to pheromones, appears to be of little relevance in *Fukomys* and other African mole-rats. The bathyergid vomeronasal organ is growth-deficient (Dennis et al., 2019; Jastrow et al., 1998) and its vomeronasal receptor repertoire is small. The latter is typical for subterranean rodents in general (Jiao et al., 2019). However, there are still structural indications for the bathyergid vomeronasal organ being functional (Dennis et al., 2019) and even if that is not the case, pheromones may also be perceived by the primary olfactory epithelium (Wang et al., 2007), so that a pheromone function for perioral compounds cannot be excluded.

### Synopsis and Conclusion

We have shown that Northern common mole-rats (as well as naked mole-rats) of both sexes possess complex and structurally derived sebaceous glands situated in their mouth corners. These perioral glands show a sex-specific secretion activity, which is higher in males and not affected by reproductive status. Conspecifics of both sexes, but particularly females, are notably responsive to male perioral secretion, which suggest them to serve in social communication. Finally, metabolic profiling revealed that the composition of volatile compounds of perioral sebum is sexually dimorphic and also allows the differentiation of female and male breeders and non-breeders.

These results suggest that perioral secretions convey important sex-specific social information in both sexes but that perioral signaling is asymmetrical, as males invest more into the quantity of perioral secretions. It appears possible that while female secretions might be exclusively perceived during close contact (i.e. facial nuzzling) the greater quantities of male secretion enable a more potent olfactory signal that might play an important role in attracting and courting mates. It is tempting to speculate that scents derived from the perioral glands are passively deposited onto the soil during tooth digging. The characteristic waxy consistency of the secretion might aid in prolonging the longevity of the scent signal (compare Scordato et al., 2007). This way, for instance, male tenants could effectively signal their presence and perhaps their physical condition in a burrow system to same-sex intruders that might challenge their position (Torrents-Ticó et al., 2018; Mynhardt et al., 2021).

The perioral secretions of *Fukomys* can be added to a long list of sexually dimorphic traits in these monogamous, cooperatively-breeding rodents, all pointing to a notable role of male intrasexual competition within their social system (Caspar et al., 2021a). Yet, it is intriguing to note that not all populations of *Fukomys* seem to express conspicuous perioral stains and that there is pronounced intrasexual variation. The physiological causes and behavioral implications of this variability will need to be clarified.

Several further new questions on olfactory signaling in common mole-rats and other bathyergids arise from this research. For instance, it remains to be determined how perioral and anal gland derived scents complement each other in the mediation of social behaviors and whether traces of perioral sebum can indeed act as lasting scent marks in burrow systems. It is to hope that such work on the conspicuous perioral secretions of social mole-rats will not only clarify how these animals effectively communicate underground, but also stimulate research on the understudied mouth corner glands of other rodent species.

## Acknowledgments

We are indebted to Radim Šumbera (České Budějovice) and Pavel Němec (Prague) for granting us access to their mole-rats. We thank Jan “Honza” Okrouhlík for practical assistance in České Budějovice and Christiane Wittmann and Patricia Gerhardt for providing us access to essential laboratory infrastructure in Essen. Kyle Finn is acknowledged for helpful discussions. KRC was supported by a Ph.D. fellowship of the German Academic Scholarship Foundation (Studienstiftung des deutschen Volkes e.V.).

## Author contributions

KRC, SB and PS designed the study; DI, KK, KRC, and SB collected behavioral and morphological data, KRC, SZ & TZ performed histology and quantified respective data, PZ conducted the GCxGC-MS experiments, KRC & PS analyzed and curated the data, KRC & PS wrote the manuscript with input from all remaining authors.

## Data availability

The paper and its accompanying supplementary information contain all data discussed in the study.

## Ethics declarations

The authors declare no competing interests.

