## Supplementary figures and images for "Perioral secretions enable complex social signaling in African mole-rats (genus *Fukomys*)"

### Supplementary Figure 1

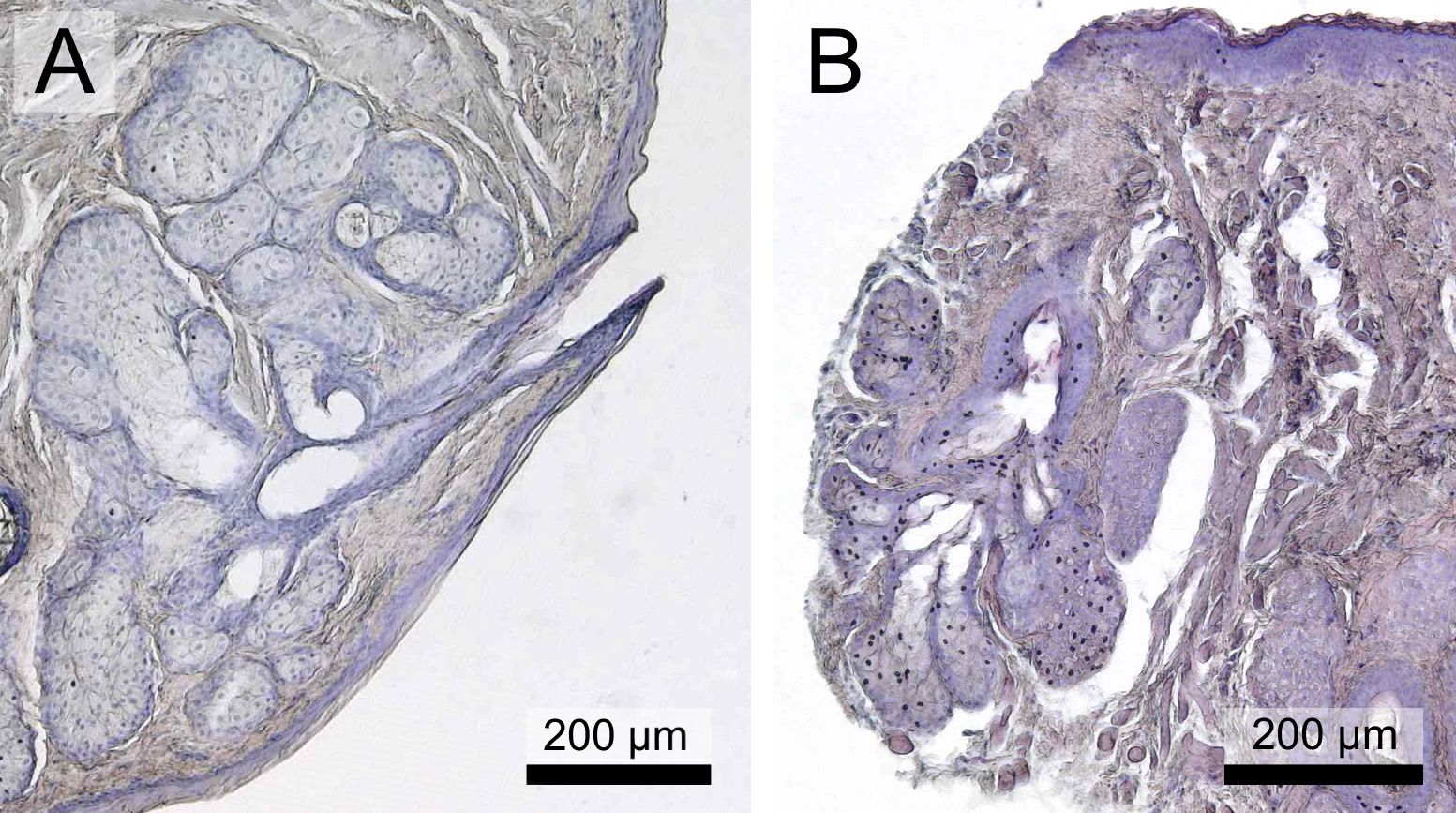

### Supplementary Figure 2

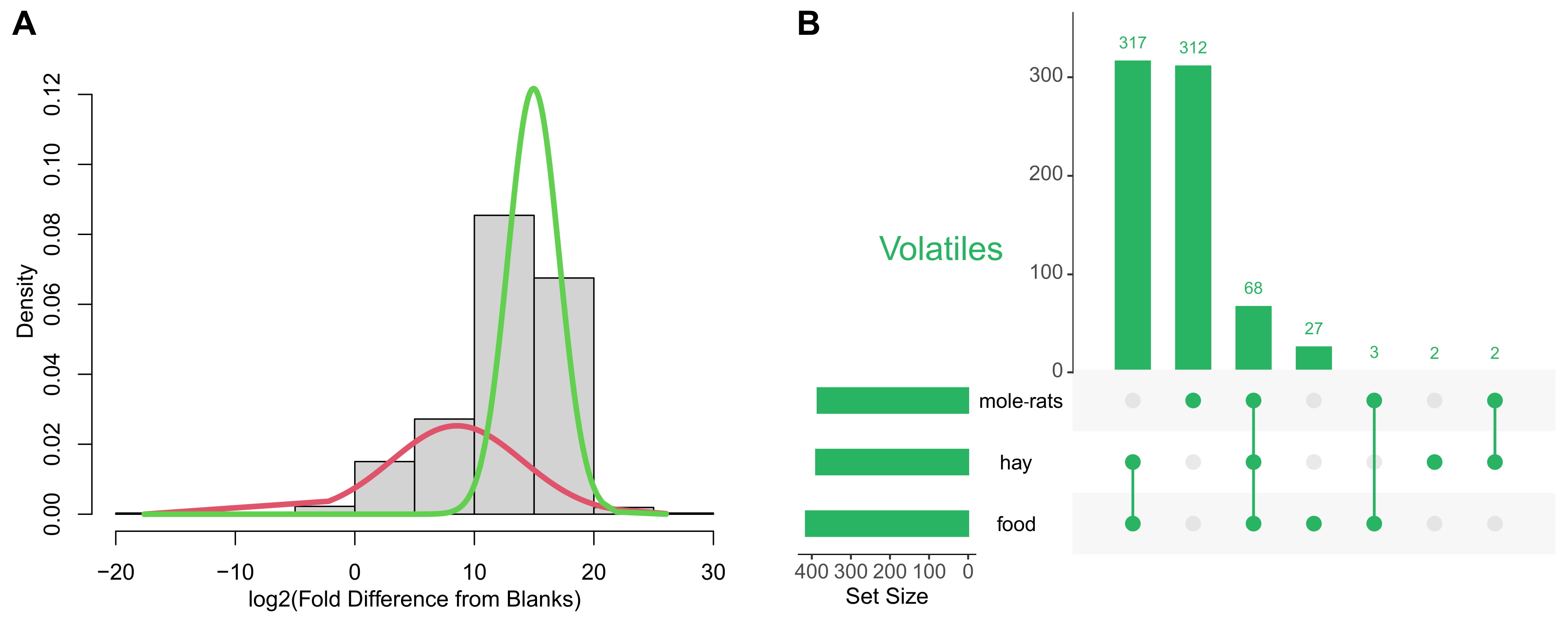
